# Arc restores juvenile plasticity in adult mouse visual cortex

**DOI:** 10.1101/130880

**Authors:** Kyle R. Jenks, Taekeun Kim, Elissa D. Pastuzyn, Hiroyuki Okuno, Andrew V. Taibi, Haruhiko Bito, Mark F. Bear, Jason D. Shepherd

**Affiliations:** Department of Neurobiology and Anatomy, University of Utah, Salt Lake City Utah, 84112; The Picower Institute for Learning and Memory, Massachusetts Institute of Technology, Cambridge Massachusetts, 02139; Medical Innovation Center, Kyoto University Graduate School of Medicine, Sakyo-ku, Kyoto 606-8507, Japan; Department of Neurochemistry, Graduate School of Medicine, The University of Tokyo, Hongo 7-3-1, Bunkyo-ku, Tokyo 113-0033, Japan

**Keywords:** Arc, plasticity, visual cortex, ocular dominance plasticity, amblyopia, critical period

## Abstract

The molecular basis for the decline in experience-dependent neural plasticity over age remains poorly understood. In visual cortex, the robust plasticity induced in juvenile mice by brief monocular deprivation (MD) during the critical period is abrogated by genetic deletion of Arc, an activity-dependent regulator of excitatory synaptic modification. Here we report that augmenting Arc expression in adult mice prolongs juvenile-like plasticity in visual cortex, as assessed by recordings of ocular dominance (OD) plasticity *in vivo*. A distinguishing characteristic of juvenile OD plasticity is the weakening of deprived-eye responses, believed to be accounted for by the mechanisms of homosynaptic long-term depression (LTD). Accordingly, we also found increased LTD in visual cortex of adult mice with augmented Arc expression, and impaired LTD in visual cortex of juvenile mice that lack Arc or have been treated *in vivo* with a protein synthesis inhibitor. Further, we found that although activity-dependent expression of *Arc* mRNA does not change with age, expression of Arc protein is maximal during the critical period and declines in adulthood. Finally, we show that acute augmentation of Arc expression in wild type adult mouse visual cortex is sufficient to restore juvenile-like plasticity. Together, our findings suggest a unifying molecular explanation for the age- and activity-dependent modulation of synaptic sensitivity to deprivation.

**Significance Statement:** Neuronal plasticity peaks early in life during critical periods and normally declines with age, but the molecular changes that underlie this decline are not fully understood. Using the mouse visual cortex as a model, we found that activity-dependent expression of the neuronal protein Arc peaks early in life, and that loss of activity-dependent Arc expression parallels loss of synaptic plasticity in the visual cortex. Genetic overexpression of Arc prolongs the critical period of visual cortex plasticity and acute viral expression of Arc in adult mice can restore juvenile-like plasticity. These findings provide a mechanism for the loss of excitatory plasticity with age, and suggest that Arc may be an exciting therapeutic target for modulation of the malleability of neuronal circuits.

## Introduction

A defining feature of early postnatal brain development is the activity-dependent winnowing of synaptic connections. This process is readily demonstrated by the response of visual cortical circuits to temporary monocular deprivation (MD) during early life. When MD is initiated during an early critical period, the synapses serving the deprived eye in visual cortex lose strength and are eliminated. Deprived-eye depression diminishes with age such that by the onset of adolescence, circuits are less vulnerable to the effects of deprivation. Understanding the molecular mechanisms that underlie the effect of age on this type of ocular dominance (OD) plasticity is one of the great challenges in neuroscience (1).

It is now well established that OD plasticity after MD occurs through synaptic plasticity of excitatory transmission, employing mechanisms that include homosynaptic long-term depression (LTD), metaplasticity and homeostatic scaling of AMPA-type glutamate receptors (2, 3). Clues into the molecular basis for the decline in juvenile plasticity have come from several diverse experimental treatments that can restore or prolong sensitivity to MD in adult animals. These include genetic manipulations that slow the maturation of cortical inhibition (4, 5), enrichment of animal housing conditions (6), increased exposure to visual stimulation (7), and enhanced modulatory neurotransmission (8). It has been suggested that a common thread connecting these varied treatments might be an increase in the ratio of excitation to inhibition (9, 10). However, it is completely unknown how, at the molecular level, general increases in cortical activity can facilitate deprivation-induced synaptic plasticity in adult visual cortex. Since the immediate early gene Arc is exquisitely sensitive to changes in cortical activity, and is essential for both OD plasticity and modification of excitatory synaptic transmission (11-13), we set out to determine whether availability of Arc limits or changes the qualities of plasticity in adults and whether up-regulating Arc levels in adult animals can restore juvenile synaptic plasticity.

## Results

### Augmentation of Arc expression in adult mouse visual cortex extends the critical period of juvenile ocular dominance plasticity

In young mice (≤ postnatal day (P) 40), the main consequence of short (3-4 days) MD is the robust loss of cortical responsiveness to stimulation of the deprived eye. A compensatory potentiation of responses to the non-deprived eye may also occur, typically observed with longer periods of MD (5-7 days) (14). Importantly, although open-eye potentiation after long duration MD is also observed in adult rodents, deprived-eye depression is only observed during the juvenile critical period in animals housed under standard laboratory conditions (15, 16). We predicted that augmenting Arc levels would prolong juvenile plasticity, as defined by closed-eye depression, past the conventional critical period in mouse visual cortex. To test this prediction we utilized a transgenic mouse line that expresses an additional allele of *Arc* tagged with mCherry in an activity-dependent manner that is driven by the *Arc* promoter in a similar manner to the previously characterized Arc-GFP Tg mouse line (17, 18) (Figure S1).

We compared the qualities of OD plasticity after short (3-4 days) MD in Arc transgenic (Arc-Tg) mice and wild-type (WT) littermate controls at P30 (juvenile) and P180 (adult) using chronic recordings of visually evoked potentials (VEPs) from binocular visual cortex contralateral to the deprived eye (Figure 1A) as previously described (11). There was no significant difference between P30 WT and Arc-Tg VEPs prior to MD and, following MD, both WT and Arc-Tg P30 mice exhibited a significant decrease in contralateral (contra; closed eye) VEP amplitudes (WT: n = 7, Baseline = 251 ± 28 μV, Post-MD =166 ± 12 μV*, p* = 0.03; Arc-Tg: n = 10, Baseline = 227 ± 21 μV, Post-MD = 159 ± 22 μV, *p* = 0.01; paired *t*-test; Figure 1B). As expected, adult P180 WT mice did not exhibit depression of contra VEP amplitude after MD, reflecting the loss of juvenile plasticity. In sharp contrast, P180 Arc-Tg mice still exhibited a significant decrease in contra VEPs (WT: n = 7, Baseline = 184 ± 19 μV, Post-MD = 183 ± 20 μV, *p* =0.9; Arc-Tg: n = 6, Baseline = 208 ± 26 μV, Post-MD = 136 ± 20 μV, *p* = 0.02; paired *t*-test; Figure 1C), comparable to the decrease observed in WT juveniles. There was a significant treatment by genotype interaction, indicating that OD plasticity differs in Arc-Tg mice compared with WT mice (repeated measures ANOVA; *p* = 0.0092).

**Figure 1.**
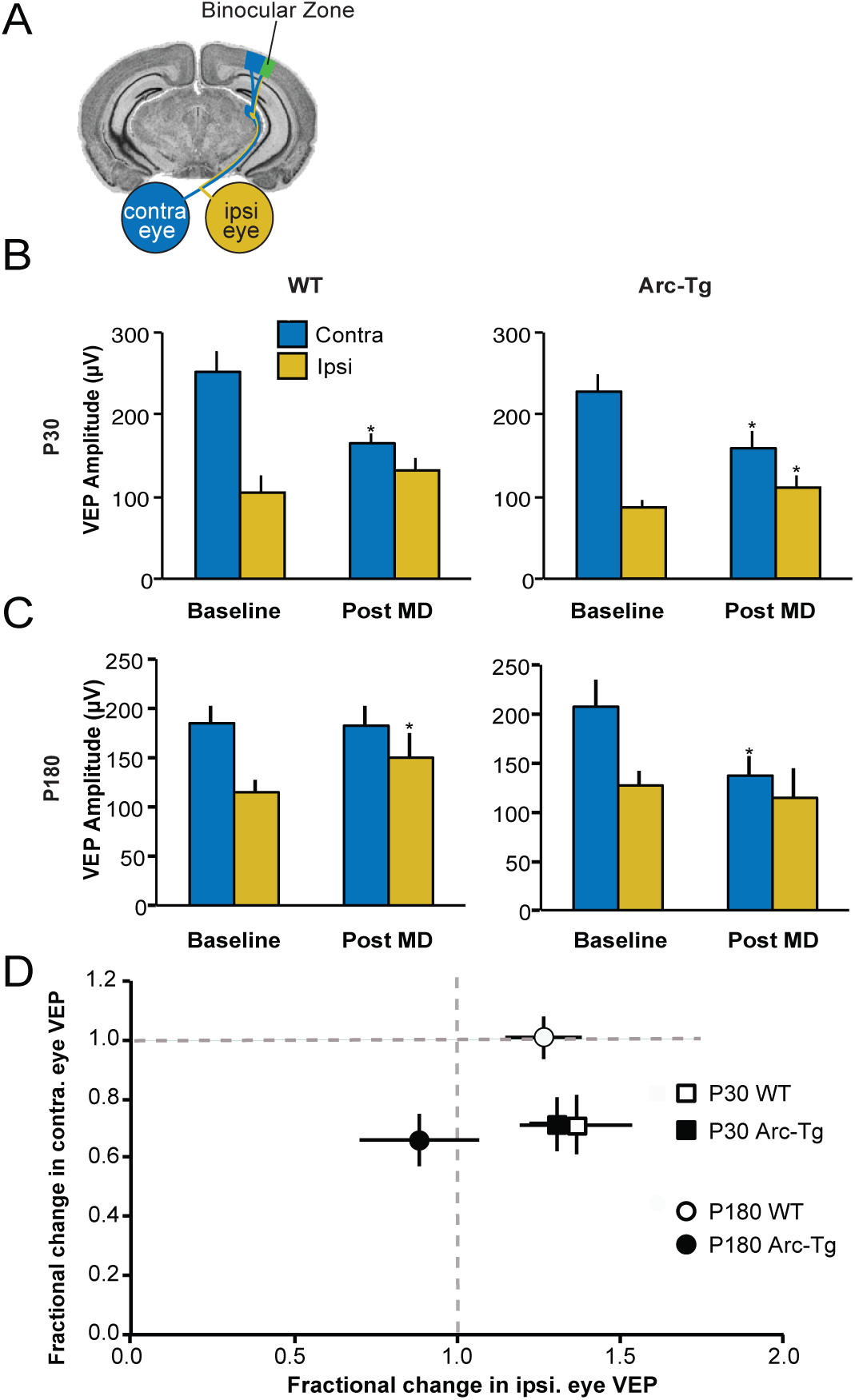
Arc-Tg mice exhibit juvenile-like OD plasticity well past the conventional critical period. (A) Schematic of recording site for VEPs in layer IV of binocular visual cortex. (B) At P30, both WT and Arc-Tg mice show a significant decrease in contralateral (closed eye/contra) VEP amplitude following MD (WT n = 7, \**p* = 0.03; Arc-Tg n = 10, \**p* = 0.01). Additionally, Arc-Tg mice exhibited a small but significant increase in ipsilateral (open eye/ipsi) VEPs (Arc-Tg *p* = 0.008). There is no significant difference between WT and Arc-Tg animals before or after MD. (C) At P180, only Arc-Tg mice exhibit a significant decrease in contra VEPs (Arc-Tg n = 6, \**p* = 0.02). (D) Plot of the fractional change in contralateral (X-axis) and ipsilateral (Y-axis) eye VEPs following MD (same data as in B and C). At P30 there is no significant difference between WT and Arc-Tg mice. However, at P180 there is a significant difference between the fractional change of WT and Arc-Tg mice following MD (*p* = 0.03). Data are represented as mean ± S.E.M.

Because the chronic VEP method enables measurements of response strength in the same mouse before and after MD, we can also analyze the qualities of the OD shift by plotting the fractional changes in response magnitude to stimulation of the deprived contra eye and the ipsi eye (19, 20). This analysis confirms that at P30, both WT and Arc-Tg mice exhibit robust and comparable levels of contralateral eye depression, and a variable potentiation of the non-deprived ipsilateral eye (Figure 1D, square symbols; WT: contralateral depression = 0.7 ± 0.1, ipsilateral potentiation = 1.4 ± 0.2; Arc-Tg: contralateral depression = 0.7 ± 0.1, ipsilateral potentiation = 1.3 ± 0.1, *p* = 0.9; MANOVA). There was, however, a significant difference in the qualities of OD plasticity in WT and Arc-Tg adult mice (Figure 1D, round symbols). In WT mice, the OD shift was accounted for entirely by ipsi eye potentiation (Fig. 1D, open circles), whereas the shift in Arc-Tg mice (Fig. 1D, filled circles) was solely due to contra eye depression (WT: contralateral depression = 1.0 ± 0.01, ipsilateral potentiation = 1.3 ± 0.1; Arc-Tg: contralateral depression = 0.7 ± 0.1, ipsilateral potentiation = 0.9 ± 0.2, *p* = 0.03; MANOVA; Figure 1D).

These data show that augmenting Arc levels in adult mice prolongs juvenile-like OD plasticity, as evidenced by deprivation-induced synaptic depression, well past the conventional critical period in mice.

### Activity-dependent Arc protein expression is high during the critical period and low in adulthood

We reasoned that if availability of Arc influences the qualities of OD plasticity, Arc expression might decline as the animal ages. In mouse visual cortex, Arc is first detected after eye-opening (∼P14) and expression steadily increases until ∼P30, corresponding to the age of peak sensitivity to MD (21). To determine whether Arc levels decline with age, WT or Arc-Tg mice were sacrificed at P30 or P180. Basal Arc expression in visual cortex is highly variable under standard housing conditions (21); therefore, we housed mice in the dark for 24 h, then either sacrificed them immediately (“dark” condition), or exposed them to light for 2 h (“light” condition) before sacrifice (n = 6/group) (22). The brain was fixed, sectioned at 30 μm on a cryostat, and immunohistochemistry (IHC) was performed for Arc protein on sections of brain containing primary visual cortex. The integrated density of Arc-expressing cells in layer IV of visual cortex was measured with the experimenter blind to genotype and age (Figure 2A). A three-way ANOVA comparing genotype (WT or Arc-Tg), age (P30 or P180), and condition (dark or light) revealed a main effect of genotype (*p* < 0.0001), age (*p* = 0.02), and condition (*p* < 0.0001), as well as a genotype x condition interaction (*p* = 0.02). *Post hoc* Student’s *t*-tests showed that in P30 mice, light significantly induced Arc expression in both WT and Arc-Tg mice (WT: light > dark; light: 4.5 ± 1.3, dark: 1 ± 0.6, *p* = 0.02; Arc-Tg: light > dark; light: 8.2 ± 1, dark: 2.7 ± 2.7, *p* = 0.002). However, Arc-Tg mice expressed significantly more Arc after light exposure than WT mice (*p* = 0.008). At P180, WT mice no longer showed detectable Arc expression, even after light exposure. Arc-Tg mice, on the other hand, exhibited significant Arc expression after light exposure (light: 7.1 ± 0.8, dark: 1.8 ± 1.2; *p* = 0.001). Furthermore, levels of light-induced Arc in P180 Arc-Tg mice were not significantly different from P30 Arc-Tg mice (*p* > 0.05), suggesting that activity-dependent expression of Arc in Arc-Tg mice does not decline with age. These data show that activity-dependent Arc protein expression significantly declines with age in WT but not in Arc-Tg mice. This loss of endogenous Arc protein over age correlates with the decline of deprived-eye depression following MD.

**Figure 2.**
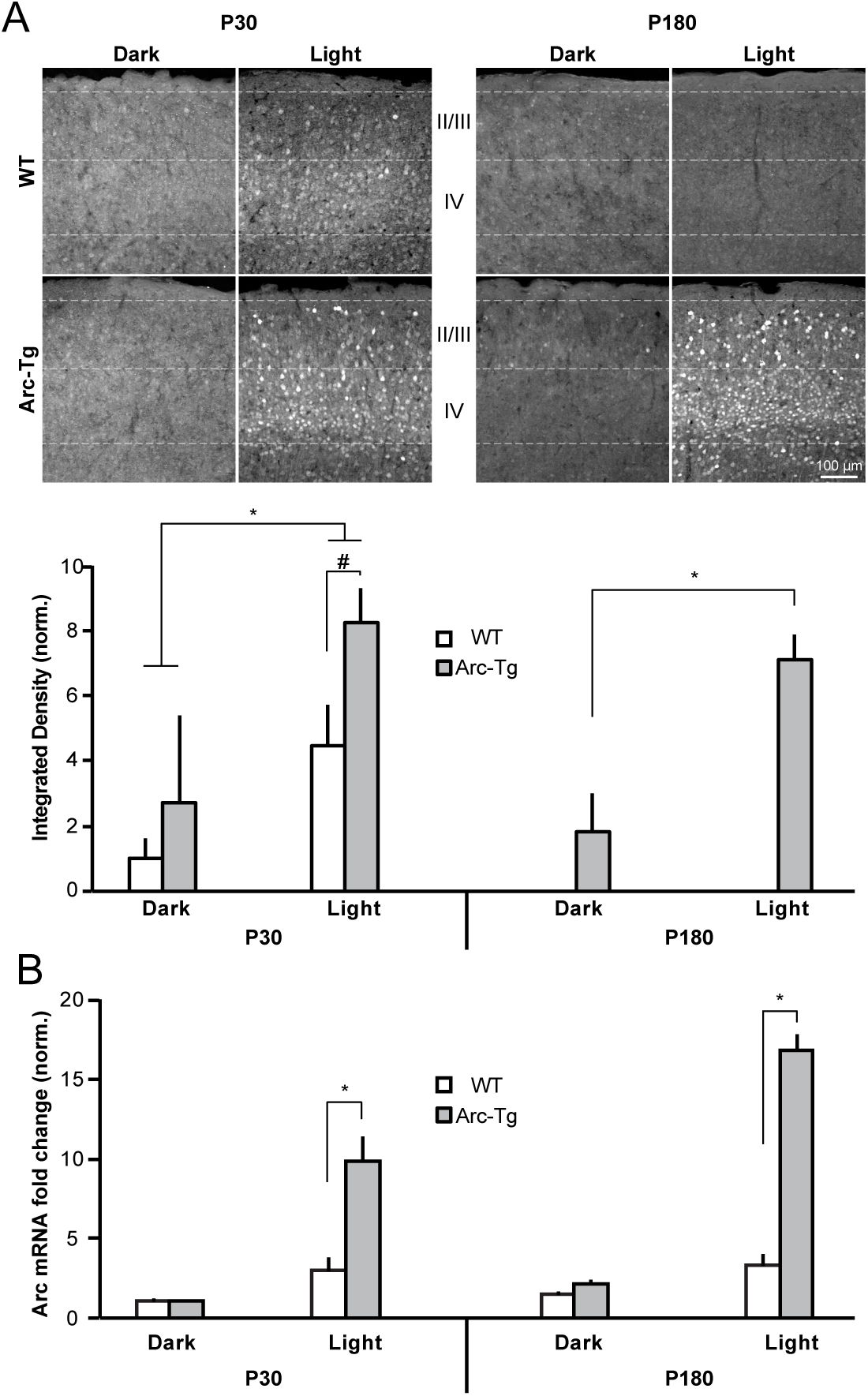
Activity-dependent Arc protein but not mRNA expression declines with age in WT mouse visual cortex but not in Arc-Tg mice. (A) Immunohistochemistry for Arc expression in layers I-IV of visual cortex after 24 h of being housed in the dark, or 24 h of dark-housing followed by 2 h of light exposure. Layer IV Arc expression is quantified in the graphs (n = 6/group). Light increased Arc expression in both WT and Arc-Tg mice at P30 (WT \**p* = 0.02; Arc-Tg *p* = 0.002), but Arc levels were higher in Arc-Tg mice (#*p* = 0.008). At P180, WT mice did not express Arc after light exposure, while Arc-Tg mice exhibited the same light-induced increase in Arc observed at P30 (\**p* = 0.001). Scale bar = 100 μm. (B) WT and Arc-Tg mice were dark housed for 24 h and then either sacrificed in the dark (“dark” condition) or exposed to light for 2 h prior to sacrifice (“light” condition). qRT-PCR was run on dissected visual cortex to quantify Arc mRNA expression. All values were first normalized to GAPDH to control for total RNA levels. Light induced Arc mRNA expression was higher in Arc-Tg mice than WT mice at both P30 and P180 (P30 \**p* < 0.0001; P180 \**p* < 0.0001). However, light-induced mRNA expression did not decrease with age in WT mice. Plotted data is normalized to P30 WT dark (n = 5 for WT Light, n = 4 for Arc-Tg light, and n = 3 for all dark groups). Data are represented as mean ± S.E.M.

Arc transcription and translation are exquisitely regulated in the brain and are finely tuned to experience and neuronal activity (12). Of particular interest, transcription and translation of Arc can be independently regulated by activity (23). We therefore sought to determine whether endogenous activity-dependent *Arc* mRNA expression also declines with age. Mice underwent dark and light exposure as described above (n = 3-5/group). The visual cortex was dissected and RT-qPCR was performed on lysates (Figure 2B). A three-way ANOVA revealed a main effect of genotype (*p* = 0.002) and condition (*p* = 0.0002) but not age. *Post hoc t*-tests showed that light-induced *Arc* mRNA expression was higher in Arc-Tg than WT mice (P30 WT: 2.9 ± 0.9, P30 Arc-Tg: 9.8 ± 1.6, *p* < 0.0001; P180 WT: 3.3 ± 0.7, P180 Arc-Tg: 16.7 ± 1.1, *p* < 0.0001). Interestingly, however, levels of activity-induced *Arc* mRNA expression did not differ with age in either genotype (*p* > 0.05). These data suggest that availability of endogenous *Arc* mRNA alone cannot fully explain the differences in Arc protein expression across the lifespan of WT mice and point to the possibility of a decrease in either activity-dependent translation or stability of endogenous Arc protein in adult visual cortex. Nevertheless, the increased expression of mRNA in the active visual cortex of Arc-Tg mice is paralleled by a proportional increase in protein.

### Augmenting Arc expression restores LTD in adult visual cortex

Deprived-eye depression occurs via mechanisms shared with LTD (3), which also diminishes with age (24). In addition to the profound deficit in OD plasticity (11), juvenile (P20-25) Arc knock-out (KO) mice also exhibit impaired layer IV LTD in visual cortex, induced in slices with low-frequency stimulation (LFS) of the white matter, as compared with WT mice that showed robust LFS LTD (WT: n = 7 slices, 4 mice 67.5 ± 5.7%; Arc KO: n = 7 slices, 5 mice 90.6 ± 4.6%; *p* < 0.001, *t*-test; Figure 3A). We therefore hypothesized that the persistence of juvenile OD plasticity in adult Arc-Tg mice was accompanied (and perhaps accounted for) by continued expression of juvenile-like LTD. To ensure expression of Arc protein in the slices, mice were exposed briefly (30 min) to an enriched environment prior to sacrifice as described previously (23). We measured LTD at P30-40, when both WT and Arc-Tg mice show comparable juvenile OD plasticity, characterized by robust deprived-eye depression after MD. At this age, LTD in WT and Arc-Tg mice was also comparable (WT: n = 9 slices, 7 mice 75.4 ± 11.6%; Arc-Tg: n = 7 slices, 6 mice 81.3 ± 7.1%; *p* > 0.5, *t*-test; Figure 3B). However, in striking agreement with the findings of juvenile levels of deprived-eye depression following MD (Figure 1), we found that LFS induced significant LTD in adult (P180-200) Arc-Tg slices but not in WT littermate slices (WT: n = 11 slices, 6 mice 102.8% ± 8.7; Arc Tg: n = 12 slices, 6 mice 74.5 ± 7.9%; Figure 3C). The difference between genotypes was significant (*p* = 0.04, *t*-test).

**Figure 3.**
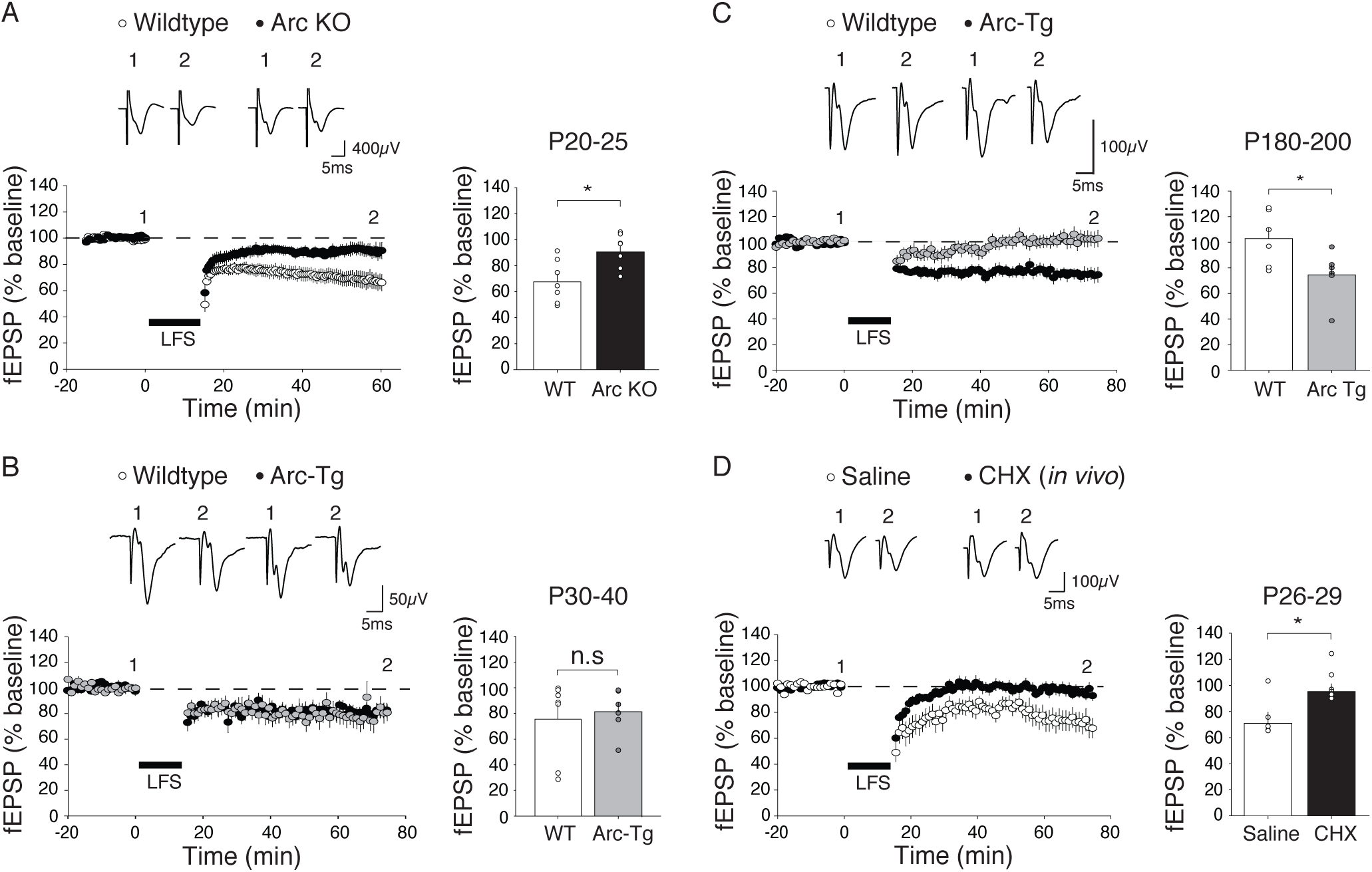
Arc and protein translation are required for LTD in layer IV of visual cortex. (A) Low frequency stimulation (LFS, 900 stimuli at 1 Hz) induces robust LTD in juvenile (P20-P25) WT but not Arc-KO slices (average of last five minutes of recordings normalized to the baseline; WT n = 4 mice; KO n = 5, \**p* < 0.001). (B) LFS induced LTD to the same degree in young (P30-40) WT and Arc-Tg slices (WT n = 7; Arc-Tg n = 6, p > 0.5). (C). LFS induces robust LTD in adult (P180-200) Arc-Tg but not WT slices (WT n = 11; Arc-Tg n = 6, \**p* = 0.04). (D) LFS induced LTD in juvenile (P25-30) visual cortex previously infused with saline but not in visual cortex infused with cycloheximide (CHX) (Saline n = 4; CHX n = 5, \**p* = 0.02). Data are represented as mean ± S.E.M.

### Inhibition of protein synthesis *in vivo* impairs LTD in juvenile visual cortex

The apparent requirement of Arc translation for deprived-eye depression may offer a partial explanation for why juvenile OD plasticity following brief MD is impaired when the visual cortex is infused locally with the protein synthesis inhibitor cycloheximide (CHX) (25). If this explanation is correct, and the mechanisms of LTD are utilized for deprived eye depression following MD, we would also expect to observe reduced LTD *ex vivo* following microinfusion of CHX into visual cortex. To test this prediction, WT visual cortex was infused *in vivo* via an osmotic minipump with CHX for four days as described (25), and then slices were prepared to study LTD. Similar to our observations in the Arc KO, there was no LTD in juvenile visual cortex after chronic inhibition of protein synthesis (saline: n = 5 slices, 4 mice, 72.4 ± 8.6%; CHX: n = 7 slices, 5 mice, 96.2 ± 5.9%*; t*-test, *p* = 0.02, Figure 3D). Together, these findings are consistent with the hypothesis that translation of Arc gates the mechanism of deprivation-induced synaptic depression in visual cortex.

### Acute expression of Arc in adult mouse visual cortex is sufficient to re-open the critical period of juvenile ocular dominance plasticity

Augmenting the availability of Arc protein throughout development and into adulthood prolongs the critical period for juvenile OD plasticity (Figure 1). However, this does not address whether restoring Arc protein expression is sufficient to re-open the critical period of OD plasticity once it has closed. To determine whether acutely increasing Arc protein in adult visual cortex is sufficient to restore juvenile-like plasticity, we expressed Arc using a lentivirus injected into visual cortex of P180 WT mice (Figure 4A). Lentivirus containing GFP-Arc or GFP was injected into layer IV of visual cortex and baseline VEP recordings were conducted one week after virus injection. Unlike the Arc-Tg mice, viral Arc over-expression is constitutively driven and not activity-dependent. Based on previous studies (26, 27), we predicted that VEP amplitude might be depressed by constitutive Arc expression since the VEP is mainly a synaptic population response that correlates with surface AMPAR expression (11). Indeed, a significant decrease in overall binocular VEP amplitude was observed compared with GFP-injected mice (GFP-injected mice: 197 ±30 μV; GFP-Arc-injected mice: 75 ± 21 μV; *p =* 0.005; Figure 4B). No deprived-eye depression was observed in GFP-injected mice following short (3-4 days) MD (GFP, normalized to baseline contra values: n = 11, contra Baseline = 1 ± 0.2, Post-MD = 0.9 ± 0.2, *p* = 0.4; paired *t*-test; Figure 4C). However, despite a reduction in baseline VEP magnitude, contra VEP responses were further reduced after MD in GFP-Arc-injected mice (GFP-Arc, normalized to baseline contra values: n = 5, contra Baseline = 1 ± 0.2, Post-MD = 0.6 ± 0.2, *p* = 0.02; Figure 4D). Further, when comparing the fractional change in contra and ipsi-eye visual responses following MD, there was a significant difference between GFP vs. GFP-Arc injected mice (GFP: contralateral depression = 0.9 ± 0.1, ipsilateral potentiation = 1.4 ± 0.1; GFP-Arc: contralateral depression = 0.5 ± 0.1, ipsilateral potentiation 1.0 ± 0.1; *p* = 0.01, MANOVA; Figure 4E). Critically, the fractional OD shift in the P180 GFP-injected mice was the same as non-injected WT P180 (*p* = 0.3; MANOVA), indicating virus injection had no effect on cortical responses or OD plasticity. Additionally, the fractional OD shift in P180 WT mice injected with GFP-Arc did not significantly differ from age matched Arc-Tg mice, indicating that acute, viral expression of Arc can restore OD plasticity to a similar degree to that achieved by transgenic augmentation of Arc throughout life (*p* = 0.6; MANOVA; Figure 4E). Intriguingly, not only was contra-eye depression observed in Arc-Tg mice but a lack of ipsi-eye potentiation was also observed, further suggesting that Arc protein levels control the qualitative aspects of OD plasticity.

**Figure 4.**
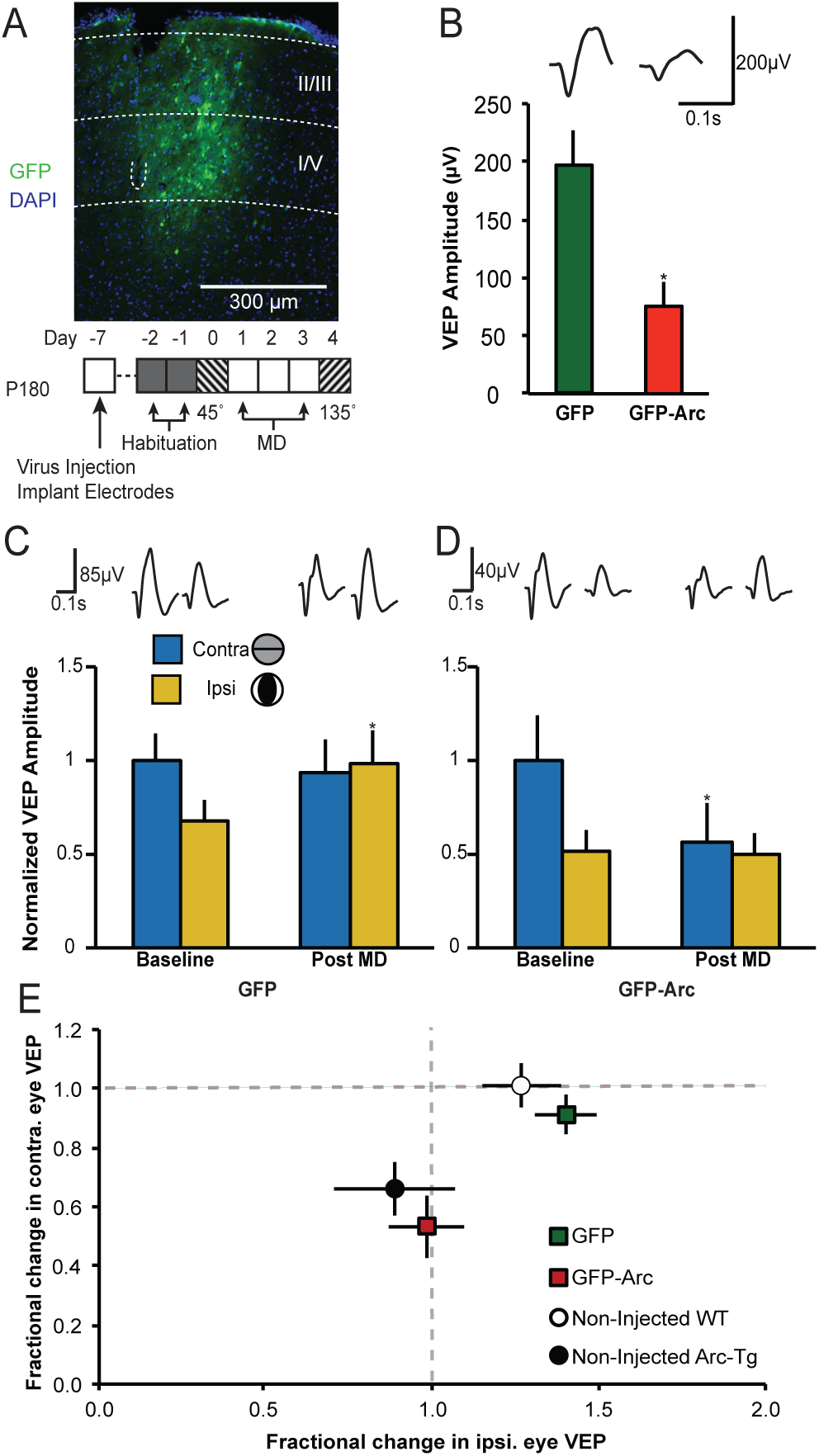
Acute Arc expression in adult mouse visual cortex is sufficient to restore juvenile OD plasticity. P180 WT mice were injected unilaterally in the visual cortex with lentivirus expressing either GFP alone or GFP-Arc. (A) Representative image of virally driven GFP expression in binocular visual cortex and timeline of the experiment. The white dashed lines demarcate the cortical layers, as well as the position of the tip of the recording electrode. (B) GFP and GFP-Arc injected P180 mice were visually stimulated prior to MD with both eyes open to record binocular baseline VEPs. GFP-Arc injected mice had significantly smaller VEPs than GFP injected mice (GFP n = 11; GFP-Arc n = 5, \**p =* 0.005). Traces represent average VEPs for GFP and GFP-Arc injected mice. (C) Data was normalized to baseline contra values. There was no significant change in Contra VEP amplitudes following MD in GFP-injected animals (*p* > 0.05), however there was a significant ipsi increase (\**p* = 0.003). (D) Data was normalized to baseline contra values. GFP-Arc injected mice exhibited significant contra depression following MD (\**p* = 0.016) and no change in ipsi responses. Averaged VEP traces are presented above the graphs. (E) Plot of the fractional change in contralateral (X-axis) and ipsilateral (Y-axis) eye VEPs following MD (same data as in C and D; Non-injected WT and Arc-Tg data from Figure 1 B). There is a significant difference between the fractional change in visual responses between GFP and GFP-Arc injected mice (*p*<0.01). GFP-injected mice exhibit the same lack of change as non-injected P180 WT mice (*p*=0.3), while GFP-Arc injected mice exhibit the same degree of change as non-injected P180 Arc-Tg mice (*p*=0.6). Data are represented as mean ± S.E.M.

These data show that acutely increasing Arc protein expression in visual cortex is sufficient to restore juvenile OD plasticity in adult visual cortex, suggesting the availability of Arc protein is sufficient to allow deprivation-induced synaptic depression in adult visual cortex.

## Discussion

Here we show that acute or chronic up-regulation of Arc protein in adult mice renders visual cortical synapses sensitive to deprived-eye depression following MD, recapitulating juvenile critical period OD plasticity. In agreement with the prevailing hypothesis that LTD mechanisms mediate deprived eye depression (3), overexpression of Arc also prolongs juvenile-like LTD in adult visual cortex. Conversely, elimination of Arc expression or inhibition of mRNA translation in juvenile visual cortex prevents both deprived-eye depression after MD *in vivo* and LTD *ex vivo*. Together, these data indicate that availability of Arc is critical for the expression of juvenile plasticity in visual cortex.

Considering the key role for Arc in determining the qualities of OD plasticity in visual cortex of juvenile animals, we predicted that the loss of deprived-eye depression after MD in adult visual cortex correlates with a lack of activity-dependent Arc expression. Indeed, we found that endogenous Arc protein expression in the active visual cortex declines with age, coincident with the loss of juvenile plasticity. Surprisingly, however, we found that activity-dependent *Arc* mRNA expression is comparable in juvenile (∼P30) and adult (∼P180) WT mouse visual cortex. This finding implies that the normal decline in Arc protein expression in active visual cortex results from a decrease in experience-dependent *Arc* translation, which can occur via mechanisms that are distinct from those regulating activity-dependent transcription (12, 23). The lack of decline in activity-dependent Arc expression in Arc-Tg mice could be due to the increase in *Arc* mRNA levels. Alternatively or in addition, the extra Arc allele in the Arc-Tg line does not contain an intron in the 3’UTR region, which may result in an increase in mRNA stability in dendrites due to a lack of nonsense mediated decay (28) and would thus potentially have a longer half-life than endogenous *Arc* mRNA. Restoration of juvenile plasticity in adult mice injected with GFP-Arc suggests that the presence of Arc protein in visual cortex is sufficient for juvenile OD plasticity.

Deprived-eye depression after MD is believed to occur via mechanisms revealed by the study of LTD in layer IV. LTD in this layer is triggered by NMDA receptor activation and expressed by internalization of AMPA receptors (29). Although NMDA receptor-dependent LTD is not affected by acute (*in vitro*) inhibition of protein synthesis (30), we discovered that chronic inhibition of protein synthesis by *in vivo* microinfusion of CHX, which has been shown to prevent deprived-eye depression (25), impairs layer IV LTD *ex vivo*. These findings are reminiscent of the recent observation that chronic, but not acute, inhibition of metabotropic glutamate receptor 5 (mGluR5) can disrupt both deprived-eye depression after MD and LTD in layer IV (19). Activity-dependent synthesis of Arc protein occurs downstream of mGluR5 activation (12, 23). Thus a simple explanation for this constellation of findings is that NMDA receptor-dependent LTD and deprived eye depression require Arc protein as a necessary cofactor, and are thus inhibited by chronic block of either mGluR5 or protein synthesis.

Decreased availability of Arc, and a consequent down-regulation of the mechanisms of LTD, also offers a simple molecular explanation for the age-dependent loss of synaptic sensitivity to visual deprivation.

Inhibition develops later than excitatory transmission in the cortex, and it has been suggested that the consequent decrease in the ratio of excitation to inhibition brings the critical period for juvenile plasticity to a close (10). We propose that decreasing the excitability of the visual cortex ultimately affects OD plasticity by preventing the activity-dependent expression of key activity-regulated plasticity proteins at the synapse that are important mediators of excitatory synaptic modification, such as Arc. Indeed, in addition to manipulations of inhibition, OD plasticity can be restored in adult rodents exposed to an enriched visual environment (6, 7), treated chronically with fluoxetine (8), or genetically engineered to express constitutively active CREB (31), manipulations that also increase Arc protein levels (32). The precise regulation of Arc expression during development, therefore, provides a potential mechanistic link between the maturation of inhibition and changes in the qualities of excitatory synaptic modification over the lifespan.

## Materials and Methods

### Animals

Lines of Tg mouse harboring the Arc-promoter mCherry-Arc transgene (mCherry-Arc/Arc) were generated as previously described (18). Further details can be found in supplementary methods section. Requests for mice should be directly addressed to H.B. or H.O. Arc-KO mice were obtained from Dr. Kuan Wang (NIH) and were previously described (22). Both male and female mice were used and the experimenter was blind to genotype in all experiments. Male C57BL/6 mice (Charles River Laboratories) at the age of P22-P25 were used for the Alzet pump implantation experiments. Male C57BL/6 mice (Jackson Laboratory) at the age of P180 were used for lentiviral VEP experiments. All procedures were approved by the Institutional Animal Care and Use Committees of Massachusetts Institute of Technology, the University of Utah, and the University of Tokyo Graduate School of Medicine, in conjunction with NIH guidelines.

### Virus production/injection

*Virus production:* Dr. Kimberly Huber generously donated FUGW lentiviral plasmids for Ubq-GFP and Ubq-GFP-Arc. Injections were carried out as previously described (33).

### *VEP recordings, slice electrophysiology* and *immunohistochemistry*

were carried out as previously described (11, 19). Detailed Methods on immunohistochemistry, quantitative RT-PCR, VEP recordings and slice electrophysiology can be found in supplementary methods.

### Statistics

ANOVA/MANOVA tests and *post hoc* Student’s *t*-tests were performed using JMP Pro software (v12; SAS Institute, Cary, NC). For slice electrophysiology experiments, *post hoc* paired *t*-tests were performed to determine the significance of changes before and after LFS, and unpaired *t*-tests were performed to test the differences between groups after LFS.

## Acknowledgements

K.R.J. was supported by the University of Utah Neuroscience training program (5T32NS076067) and an NIH NRSA F31 (MH112326). E.D.P. was supported by a developmental biology training grant at the University of Utah (5T32HD00749117). This work was funded by AMED-CREST (H.B.), KAKENHI grants from JSPS (H.O., H.B.), a grant-in-aid from MHLW-Japan (H.O., H.B), the NIH (R00-NS076364, J.D.S), the Howard Hughes Medical Institute, and the Picower Institute Innovation Fund (M.F.B.). We thank Dr. Roger Y. Tsien (University of California San Diego) for the gift of the mCherry plasmid. We thank Dr. Kimberly Huber (University of Texas Southwestern Medical Center) for the FUGW CMV-GFP and CMV-Arc-GFP plasmids and Dr. Kuan Wang (NIH) for the Arc KO mouse line.

## Contributions

E.D.P. performed the immunohistochemistry experiments. T.K. and J.D.S. performed slice electrophysiology experiments, T.K. performed *in vivo* infusion experiments, and K.R.J. and J.D.S performed *in vivo* VEP recordings. K.R.J prepared and injected lentivirus *in vivo*. A.V.T. conducted RT-qPCR experiments. H.O. and H.B. generated the Arc-Tg mouse line. E.D.P., T.K., K.R.J., A.V.T. and J.D.S. performed data analysis. M.F.B. and J.D.S wrote the manuscript; conceived, designed and directed the study.

## Figure Legends

**Figure S1.**
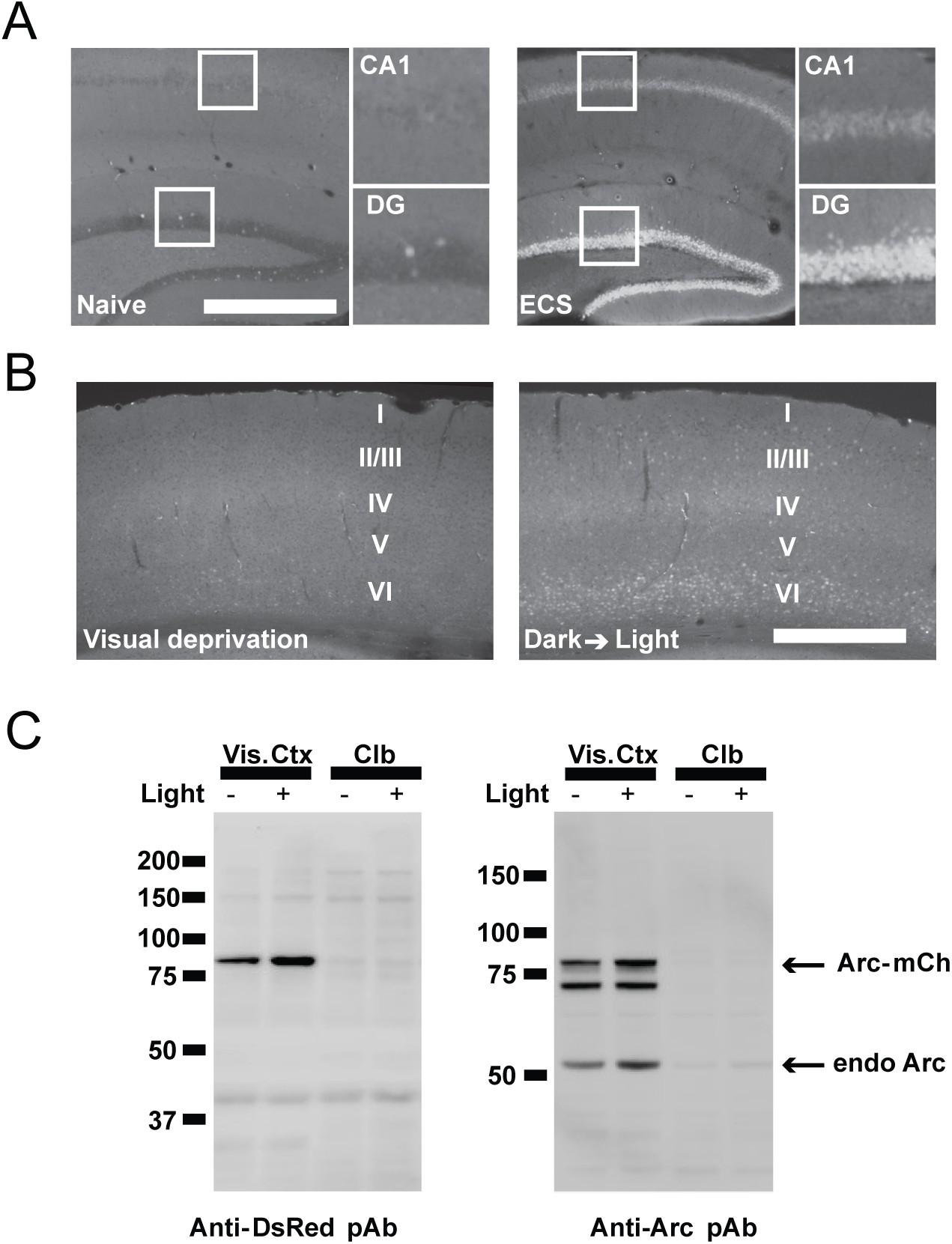

